# Phenotypic Plasticity Drives Seasonal Thermal Tolerance in a Baltic Copepod

**DOI:** 10.1101/2023.07.31.551281

**Authors:** Alexandra Hahn, Reid S. Brennan

**Affiliations:** Marine Evolutionary Ecology, GEOMAR Helmholtz Center for Ocean Research Kiel, Düsternbrooker Weg 20, Kiel, 24105, Germany

**Author notes:** Corresponding author; AH RB.

**Keywords:** CT*_max_*, Seasonality, *Acartia*

## Abstract

Seasonal changes in environmental conditions require substantial physiological responses for population persistence. Phenotypic plasticity is a common mechanism to tolerate these changes, but for organisms with short generation times rapid adaptation may also be a contributing factor. Here, we aimed to disentangle the impacts of adaptation from phenotypic plasticity on thermal tolerance of the calanoid copepod *Acartia hudsonica* collected throughout spring and summer of a single year. We used a common garden (11 °C and 18 °C) design to determine the relative impacts of plasticity versus adaptation. *Acartia hudsonica* were collected from five time points across the season and thermal tolerance was determined using critical thermal maximum (CT*_max_*) followed by additional measurements after one generation of common garden. As sea surface temperature increased through the season, field collected individuals showed corresponding increases in thermal tolerance but decreases in body size. Despite different thermal tolerances of wild collections, common garden animals did not differ in CT*_max_* within thermal treatments. Instead, there was evidence of phenotypic plasticity where higher temperatures were tolerated by the 18 °C versus the 11 °C treatment animals across all collections. Acclimation also had significant effects on body size, with higher temperatures resulting in smaller individuals, consistent with the temperature size rule. Therefore, the differences in thermal tolerance and body size observed in field collected *A. hudsonica* were likely driven by plasticity rather than adaptation. However, the observed decrease in body size suggests that nutrient availability and ecosystem functioning could be impacted if temperatures consistently increase with no change in copepod abundance. This is the first record of *A. hudsonica* in the Baltic Sea known to the authors.

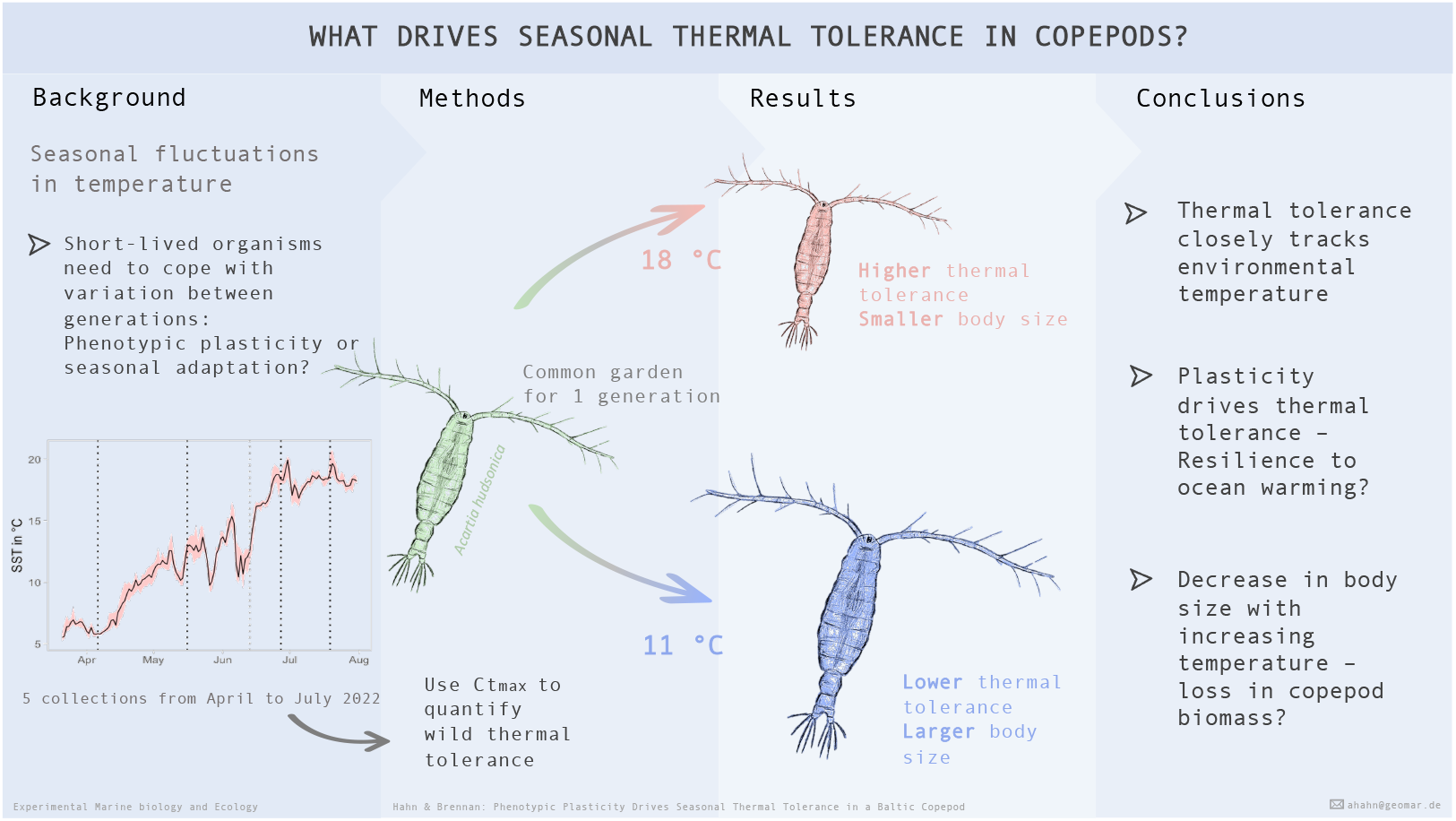
Graphical Abstract.

**Highlights:** **Phenotypic Plasticity Drives Seasonal Thermal Tolerance in a Baltic Copepod**

Alexandra Hahn, Reid S. Brennan

- *Acartia hudsonica* shows strong seasonality in thermal tolerance.
- The observed seasonal differences in CT*_max_* are driven by phenotypic plasticity not adaptation.
- Body size in *A. hudsonica* is negatively correlated to environmental and developmental temperature.
- This is the first record of *A. hudsonica* in the Baltic Sea known to the authors.

## 1. Introduction

Environmental variation is ubiquitous across habitats and organisms are able to respond to and tolerate this variation in multiple ways. When variation is both predictable and experienced within the lifespan of an individual, it is expected that plasticity will evolve (Pereira et al., 2017; Bitter et al., 2021). Conversely, if environmental variation is unpredictable or at timescales longer than generation time, plasticity is unlikely to evolve but selection should act with each environmental fluctuation. In this latter case, the resulting changes in selective pressure through time can lead to fluctuating selection, which can not only drive rapid adaptation but also contribute to the maintenance of genetic diversity in populations (Bergland et al., 2014). One of the major drivers of fluctuating selection in the wild is seasonal change. In this scenario, the environmental changes that occur within a year result in divergent selective pressures at different temporal periods. For example, summer months may favor warm tolerant genotypes while cooler spring or fall temperatures may favor genotypes that have higher performance at low temperature. While these changes are relatively consistent on a yearly basis, the temporal scale is beyond many organisms’ lifespan, which can lead to adaptation to different periods within the fluctuating seasonal environmental change. This phenomenon has been observed in diverse species and ecosystems, including marine copepods (Hairston and Dillon, 1990), *Lonchopterid* flies (Niklasson et al., 2004), dandelion (Vavrek et al., 1996), two-spotted ladybugs (Brakefield, 1985), and swallows (Brown et al., 2013), among others (Siepielski et al., 2009; Bell, 2010). There is also evidence for fluctuating selection at the genomic level: in *Drosophila* fruit flies genome-wide allele frequencies consistently and repeatedly shift between seasons due to selection, helping to maintain genetic variation within populations (Johnson et al., 2023). Thus, adaptive responses to seasonal change may be a common phenomenon across taxa with short generation times.

In addition to contributing to the maintenance of genetic variation within populations, the mechanisms underlying rapid seasonal adaptation can help shed light on how populations may respond to ongoing anthropogenic driven global change. For example, many species experience yearly temperature changes with an amplitude greater than those predicted under global warming (Bujan et al., 2020) and across terrestrial and aquatic ectotherms, populations routinely achieve increased thermal tolerance in warmer seasons (Hopkin et al., 2006; Bujan et al., 2020). This suggests that mechanisms enabling seasonal responses, such as plasticity and adaptation, might similarly drive resilience to global warming.

Copepods provide an ideal model to understand population level responses to fluctuating selection across seasons. These organisms are shortlived with generation times of a few weeks leading to multiple generations per year, each subjected to a unique thermal regime. Previous work on thermal tolerance of copepods has found contributions of both plasticity and adaptation. For instance, rearing temperature drives plastic responses and significantly influences thermal tolerance (González, 1974), egg production (Holste and Peck, 2006) and growth (Sasaki and Dam, 2020). Temperature also affects adult body size with warmer temperatures leading to smaller individuals (Viitasalo et al., 1995; Sasaki et al., 2019), consistent with the temperature-size rule (Atkinson, 1994). Conversely, there is ample evidence for local adaptation to temperature (Lonsdale and Levinton, 1985; Pereira et al., 2017; Karlsson and Winder, 2020) as well experimental evolution studies showing adaptive responses to high temperature after only a few generations (Dam et al., 2021; Brennan et al., 2022b). Finally, planktonic copepods are an integral part of marine food webs and function as an essential link between primary production and higher trophic levels (Turner, 2004; Dzierzbicka-Głowacka et al., 2019). Therefore, our understanding of drivers of copepod responses to temperature change has important implications for the resilience of marine ecosystems as a whole.

In this study, we focus on the calanoid copepod *Acartia hudsonica* (Pinhey, 1926). *Acartia* copepods are among the most-studied copepod genera, in part due to their world-wide distribution and high abundance, making them foundational to marine and coastal ecosystems (Walter and Boxshall, 2023). In the Baltic Sea, this group is one of the dominant zooplankton and critical to local ecosystems (Diekmann et al., 2012; Dzierzbicka-Głowacka et al., 2019). *Acartia hudsonica*, specifically, is a cold adapted species that is generally abundant in winter and spring months out-competing more warm adapted congeners at low temperatures. The species can tolerate a broad temperature range from at least 4 to 18 °C (Sullivan and McManus, 1986). For populations native to the Eastern United States, *A. hudsonica* produces resting eggs at temperatures > 16 °C and abundances strongly decline in summer (Sullivan and McManus, 1986).

Here, we seek to disentangle the impacts of adaptation from plasticity in thermal tolerance of *A. hudsonica* collected throughout spring and summer of 2022. We hypothesized that the thermal tolerance of wild collected individuals would closely follow the environmental temperature. Further, because developmental temperature strongly impacts copepod body size (Horne et al., 2016), we predicted that body size would decrease as temperature increased. We used common garden conditions at two different temperatures to determine if the observed thermal tolerance and body size shifts between wild individuals were driven by adaptation or plasticity. Together, these results help to reveal the underlying mechanisms driving seasonal thermal tolerance of *A. hudsonica* and provide insight into how this species may respond to warming conditions in the future.

## 2. Material & Methods

### 2.1. Sampling and cultures

All samples were collected by 100 µm WP2 net on board the research vessel *Polarfuchs* in Kiel Fjord (54°19’50”N 10°09’20”E). Live samples were stored at collection temperature until processing. The experiment included five sampling dates (referred to as collections) from April 2022 to July 2022 that spanned SST from 5.81 °C to 19.16 °C (Table 1, Hiebenthal et al. unpublished data).

**Table 1:** Overview of sampling dates and corresponding SST averages.

| Collection name | Collection date | Daily mean | 2-week mean |
| --- | --- | --- | --- |
| Collection 1 | 06 Apr 2022 | 5.81 ±0.05 °C | 6.36 ±0.47 °C |
| Collection 2 | 16 May 2022 | 12.66 ±0.53 °C | 11.44 ±0.84 °C |
| Collection 3 | 13 Jun 2022 | 12.37 ±0.33 °C | 12.86 ±1.52 °C |
| Collection 4 | 27 Jun 2022 | 18.35 ±0.13 °C | 16.55 ±1.83 °C |
| Collection 5 | 19 Jul 2022 | 19.16 ±0.79 °C | 18.11 ±0.68 °C |

Table 1: Overview over sampling dates and corresponding SST average.
| Collection name | Collection date | Daily mean | 2-week mean |
| --- | --- | --- | --- |
| Collection 1 | 06 Apr 2022 | $5.81 \pm 0.05$ °C | $6.36 \pm 0.47$ °C |
| Collection 2 | 16 May 2022 | $12.66 \pm 0.53$ °C | $11.44 \pm 0.84$ °C |
| Collection 3 | 13 Jun 2022 | $12.37 \pm 0.33$ °C | $12.86 \pm 1.52$ °C |
| Collection 4 | 27 Jun 2022 | $18.35 \pm 0.13$ °C | $16.55 \pm 1.83$ °C |
| Collection 5 | 19 Jul 2022 | $19.16 \pm 0.79$ °C | $18.11 \pm 0.68$ °C |

For all sampling dates, approximately 440 adult animals were sorted and split into two 6 L culture buckets with air supply and held at their collection temperature. Over the following two days, CT*_max_* assays were run on the wild-caught copepods. After the initial assays, the cultures were moved to a cold (11 °C) or warm (18 °C) culture room. No major fluctuations in temperature occurred throughout the experiment. All cultures were kept on a 12:12 light regime at a common salinity of 15 and were allowed to reproduce, with water changes approximately every 7 days. Feeding was *ad libitum* with *Rhodomonas sp.* and *Isochrysis galbana* (Holste and Peck, 2006; Ismar et al., 2008; Mahjoub et al., 2014).

After the initial culture establishment at treatment temperatures (2-4 days), the cultures were filtered through a nested 200 µm and 50 µm mesh sieve. Adults were retained on the 200 µm mesh and kept for further culturing. Offspring, eggs and nauplii, of the parental generation were retained on the 50 µm mesh and placed in a new culture bucket to start the F1 generation. Development was monitored to catch the onset of maturation. Once the F1 generation reached adulthood, CT*_max_* assays were repeated. Collection 3 collapsed before reaching F1. Therefore, this collection is excluded from further analysis.

### 2.2. Temperature assays

CT*_max_* was used as a proxy for thermal tolerance. CT*_max_* is defined as the temperature at which locomotion is affected in a way that prevents the individual to move away from harmful conditions, eventually resulting in death (Cowles and Bogert, 1944). In this study, CT*_max_* was the temperature at which the individuals showed no visible response to a stimulus (details below). For each collection and treatment the CT*_max_* of twenty males and twenty females was quantified. Individuals were sorted under temperaturecontrolled conditions and placed in 12 ml glass tubes filled with 5 ml of filtered seawater. Ten individuals, five males and five females were simultaneously run per trial. The starting temperature of the experimental tank matched the wild collection or culturing temperature. To minimize bias, an assistant placed the individuals in random order in the experimental tank, leaving the experimenter blinded to the animals’ sex until after the experiment. Heating was monitored with a thermometer (PCE-HPT1, PCE instruments, Menschede, Germany) placed into an additional glass tube filled with filtered seawater. The animals were given a 30-minute acclimation period before heating was started. With a 300 W and 500 W heater, heating was consistent at *∼* 0.2 °C/min (Fig. S.1). Once 22.5 °C was reached, the 300 W heater was removed, slowing the ramping temperature to *∼* 0.1 °C/min to facilitate monitoring the individuals. Throughout the experiment, the experimenter was blinded to the exact temperature. Animals were continuously monitored and when movement ceased, gentle pipetting was used to trigger a reaction. If still no movements occurred, the corresponding temperature was considered CT*_max_*.

After the experiment, the sex for all animals was confirmed and the individuals were preserved in 95% ethanol and later photographed using a Nikon imaging microscope and Nikon imaging software (NIS-Elements v. 5.20.00). All images were obtained using the same magnification to ensure consistency. From the pictures, the prosome length was measured for each copepod using ImageJ (Schneider et al., 2012). Three measurements were obtained per individual and averaged in the later analysis. While formalin is usually used to preserve plankton samples for length analysis (Connolly et al., 2017; Aguilera et al., 2020), previous work on zooplankton has shown no effect of ethanol preservation on body size (Black and Dodson, 2003) and any size effect would be consistent across the experiment. Further, ethanol preservation allows for downstream genetic analyses, which is essential when dealing with cryptic copepod species.

### 2.3. Genotyping

To confirm species identity, 4-15 copepods per collection were genotyped using the mtCOI region (Table S.1). For the DNA extraction, ethanol-preserved copepods were rinsed with ultra-clean water and rehydrated for one hour. Individual copepods were transferred into 100 µl of 5% Chelex solution, incubated for 20 minutes at 95 °C in a water bath, then centrifuged for 5 minutes at 8000 rpm. PCR reactions were conducted in 20 µl volume with 9.8 µl ultra clean water, 2 µl dNTPs, 2 µl Buffer, 2 µl LCO1490 forward primer or HCO2198 reverse primer (5mM, Folmer et al. (1994)), 0.2 µl DreamTaq DNA polymerase (Thermo Fisher Scientific Inc., Massachusetts, US), and 2 µl of DNA.

Amplification conditions were: 3 min of denaturation at 94 °C followed by 33 cycles of 45 s denaturation at 94 °C, 45 s annealing at 48 °C, 60 s extension at 72 °C and a final extension at 72 °C for 7 min. Samples were sequenced on a Sanger sequencing platform by Eurofins Deutschland (Germany). Sequences were checked and aligned to generate consensus sequences using CodonCodeAligner (CODONCODE, 2010). The *Acartia* genus consists of multiple cryptic species making species identification difficult. Therefore, we followed Figueroa et al. (2020) and used a bayesian approach to determine species and clade for all samples. Using known samples from Figueroa et al. (2020), we aligned all raw reads with MUSCLE (Edgar, 2004), converted to NEXUS format in R using APE (Paradis and Schliep, 2019), and used MrBayes to build a phylogenetic tree with *Acartia dana* as the outgroup. Tree plots were made using *ggtree* (Yu et al., 2017). Since thermal tolerance measurements showed outliers in collections 4 and 5, 38 experimental outliers and additional trial animals from those collections were genotyped.

### 2.4. Data analysis and statistics

Data manipulation, visualization, and statistics were conducted in R version 4.2.2 (R Core Team, 2022).

To understand the factors influencing CT*_max_* and length we used generalized linear models. Model selection was done by comparing model fit and Akaike’s information criterion (AIC) as well as considering biological relevance. The models included main effects of collection number (1-5), treatment (wild, warm, cold), sex, length, and the interaction of treatment and collection. Random effects of vial number, tank used for the trial, and time of day did not improve model fit and were therefore excluded from the final models. A similar model was used to determine the factors influencing copepod length. Here, we included main effects of developmental temperature and sex. Pairwise post-hoc testing was done comparing the model means using the *emmeans* package (Lenth, 2023).

## 3. Results

The sampling days spanned SST values from 5.81 °C to 19.16 °C (Fig. 1A; see Table 1 for all values and standard deviations). The thermal tolerance of wild-caught individuals mirrored the environmental temperature (Fig. 1B). CT*_max_* values were lowest in Collection 1 (25.9 °C *±*1.4 °C), where the temperature over the two weeks prior to sampling averaged 6.36 °C *±*0.47 °C. CT*_max_* then increased throughout the collections (Col-2: 27.9 °C *±*0.7 °C; Col-3: 27.9 °C *±*1.0 °C; Col-4: 29.1 °C *±*0.6 °C), reaching its maximum at Collection 5 (29.8 °C *±*2.0 °C), where the two-week SST average was 18.11 °C *±*0.68 °C (Spearman’s *ρ* = 0.758, p < 0.001). The prosome length of wild individuals showed an opposite trend, with size decreasing as SST increased (Fig 1C, Spearman’s *ρ* = –0.572, p < 0.001). Two week averages were chosen to characterize SST as this is the approximate the amount of time it takes for copepods to mature from egg to adult and can therefore be considered developmental temperature; the two-week average showed a similar trend to daily mean temperatures (Fig. S.2).

**Figure 1:**
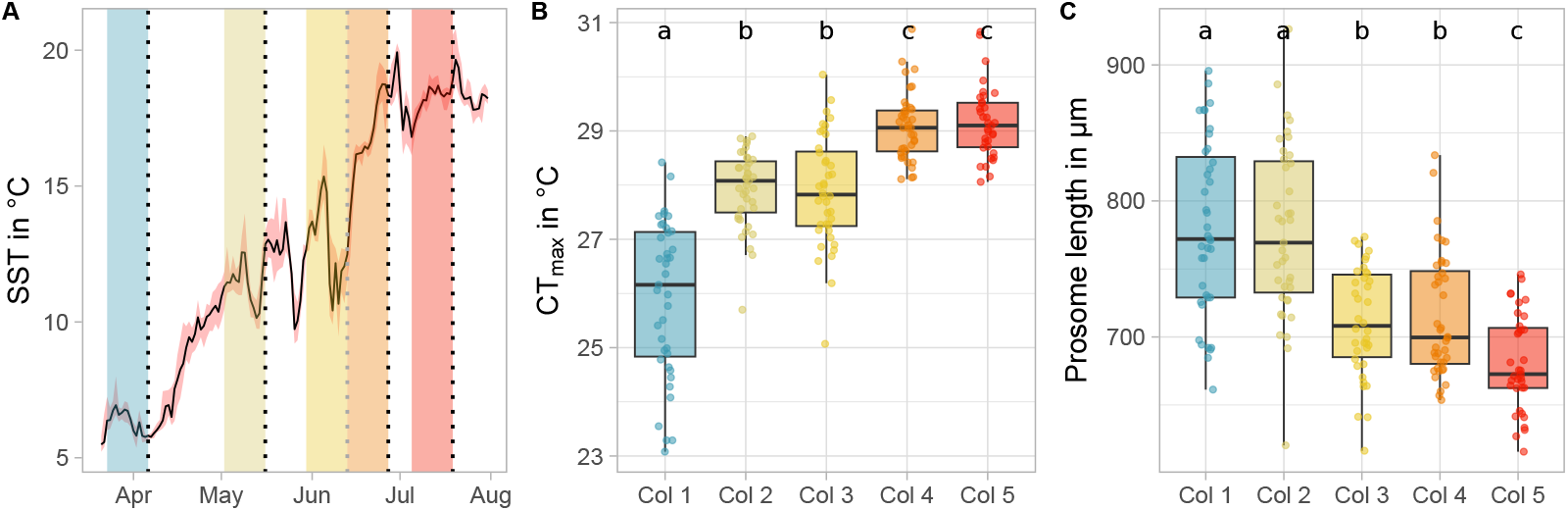
Thermal tolerance follows the seasonal changes in temperature. (A) Sea surface temperature at the collection site where sampling points are indicated by dotted lines and shaded boxes indicate the two-week period prior to sampling. (B) Critical thermal maxima and (C) mean prosome length for wild collected individuals. Boxplot colors correspond to sampling date and compact letters show the results from post-hoc tests where shared letters indicate no significant difference between measurements.

Following the common garden, there was a significant effect of treatment, collection and the interaction between treatment and collection on CT*_max_* (Fig. 2A, p *<* 0.001 for all, see Table S.3) for detailed report). While CT*_max_* of the wild individuals varied depending on the SST around sampling, the thermal tolerance measurements within the cold and warm treatment were similar across collections. The weighted model means for all warm treatments did not significantly differ from each other (col-1: 28.7 °C *±*0.5 °C, col-2: 28.6 °C *±*0.5 °C, col-4: 29.0 °C *±*0.6 °C, p *≥* 0.578) and were comparable to the wild collections 4 and 5 (29.1 °C *±*0.6 °C and 29.2 °C *±*0.7 °C respectively, p *≥* 0.153) where the mean temperature resembled the warm treatment (warm treatment 18 °C, wild SST 16.55 °C *±*1.83 °C, and 18.11 °C *±*0.68 °C respectively). For the cold treatment, weighted means did not differ significantly (col-1: 27.7 °C *±*0.5 °C, col-4: 28.0 °C *±*0.7 °C, col-5: 27.8 °C *±*0.6 °C, p *≥* 0.789) and were comparable to the thermal tolerance of the wild collection 2 (27.9 °C *±*0.7 °C, p *≥* 0.992) where the developmental temperature was similar (cold treatment 11 °C, wild SST 11.44 °C *±*0.84 °C). The exception was the cold treatment of collection 2, which was significantly lower than the other cold collections (p *<* 0.001). Here, CT*_max_* was comparable to the mean of the wild individuals within collection 1 (col-1: 25.9 °C *±*1.4 °C, col2: 26.2 *±*1.2 °C, p = 0.965) despite the treatment temperatures differing by *∼* 4 °C. The effect of treatment, collection on prosome length and the interaction of both terms were significant (Fig. 2B, p < 0.001 for all, see Table S.3). However, post-hoc testing revealed that there were no clear similarities within treatments, suggesting a more complicated connection between prosome length, seasonality and common garden temperature.

**Figure 2:**
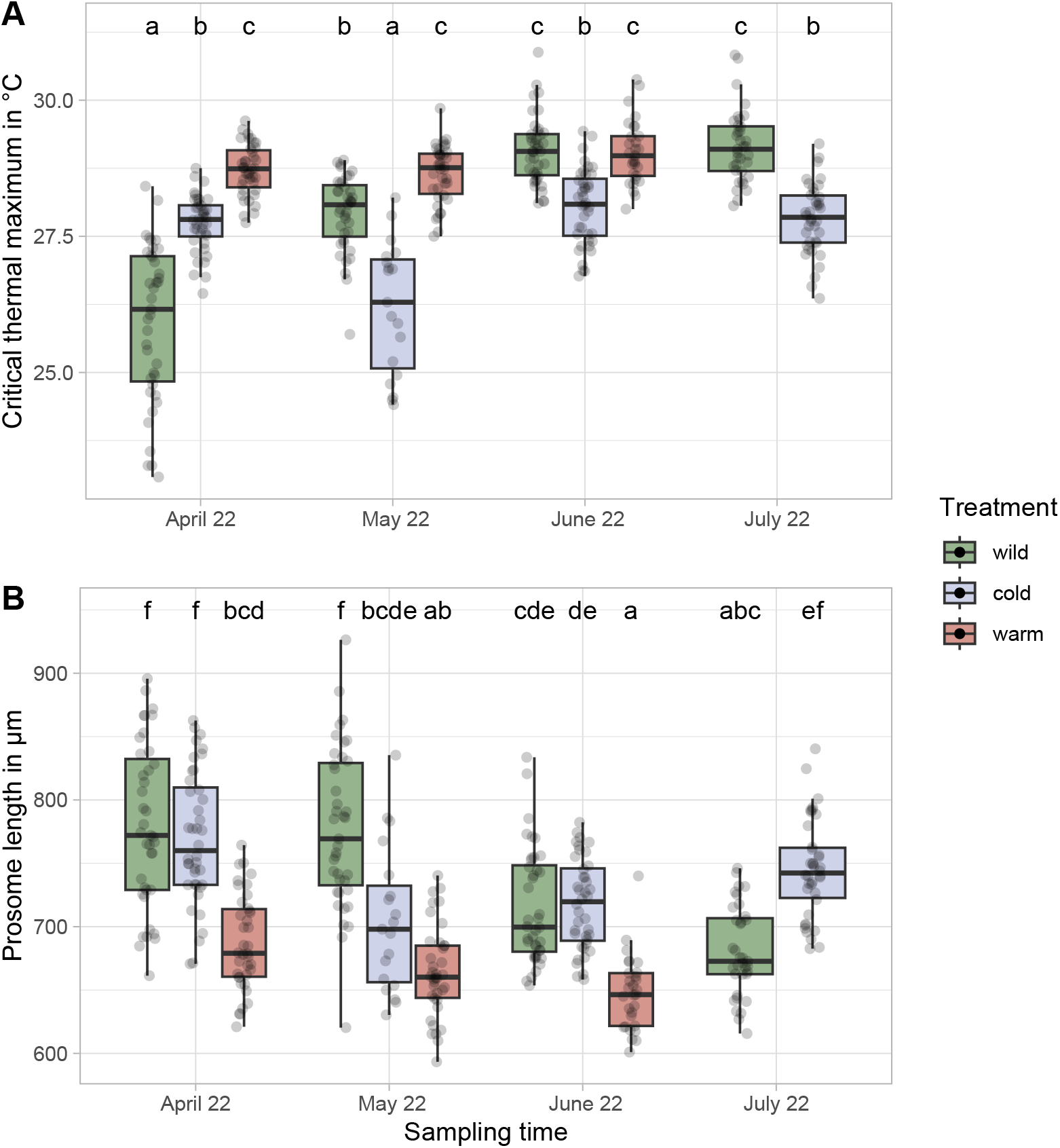
Phenotyping results following common garden for. (A) critical thermal maxima and (B) prosome length. Colors of boxes correspond to treatment where “wild” are field collected animals, “cold” are F1 animals at 11°C, “warm” are F1 animals at 18°C. Compact letters show the results from post-hoc tests where shared letters indicate no significant difference between measurements.

There was a strong plastic effect of developmental temperature on thermal performance and prosome length (p < 0.001, Fig. S.4). The reaction norms for treatment effect on CT*_max_* showed a positive effect of treatment temperature (Fig. 3A). There was a significant interaction between collections that was driven by the outlier col-2 and when col-2 was removed there was no interaction (with col-2: p, 0.001, without col-2: p = 0.897). Prosome length was negatively affected by treatment temperature (Fig. 3B). Again, a significant interaction was driven by col-2, that was not present when removing the outlier collection (with col-2: p = 0.019, without col-2: p = 0.372). The similar slopes of reaction norms and the absence of an interaction between treatment and collection support the presence of plasticity.

**Figure 3:**
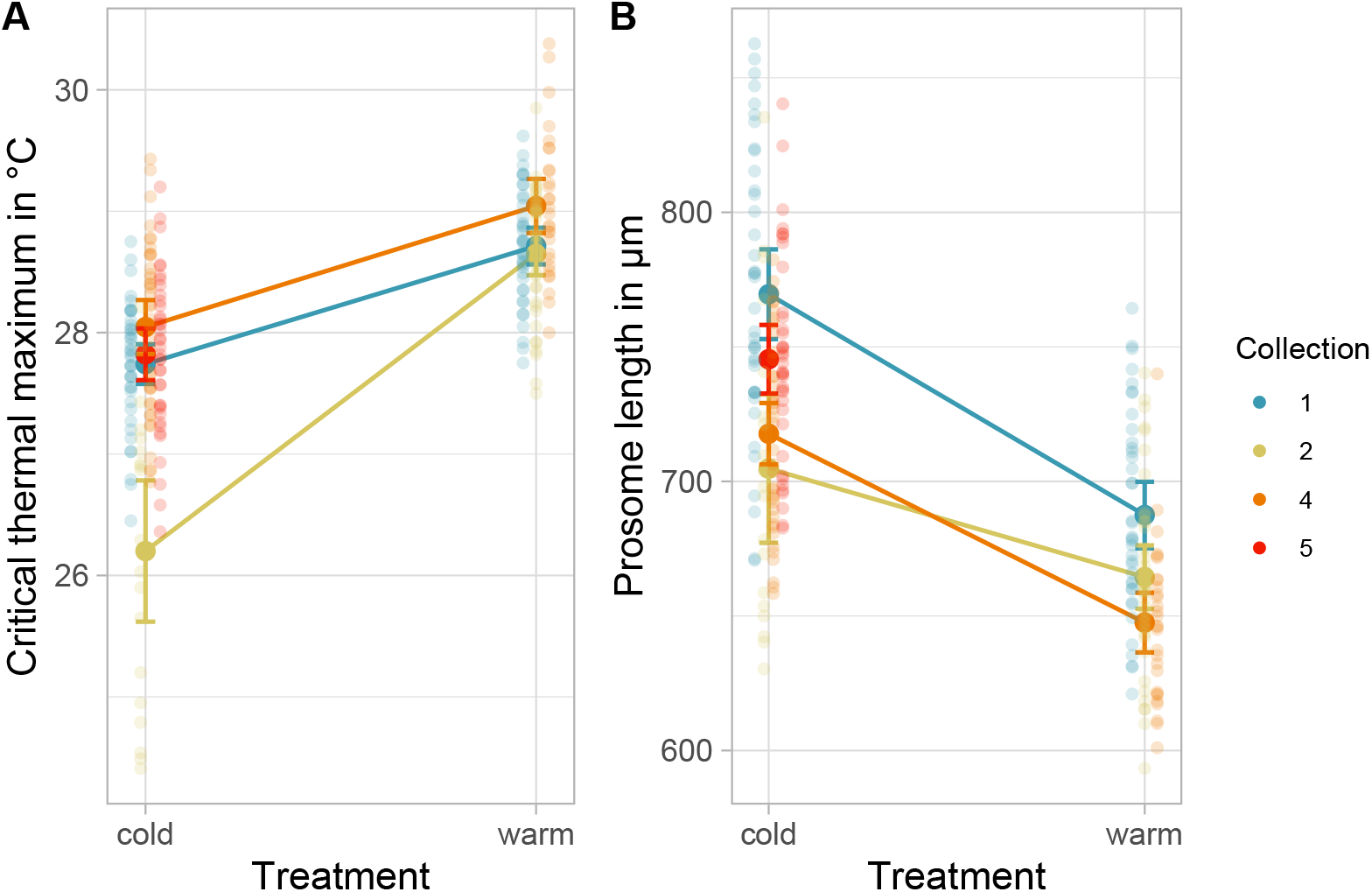
Reaction norms for treatment effect on. (A) Critical thermal maximum; (B) Prosome length; bold points indicate mean values, error bars indicate 0.95 % confidence interval.

Furthermore, there was an increase of CT*_max_* with increasing developmental temperature (Spearman’s *ρ* = 0.718, p < 0.001). Conversely, prosome length was negatively correlated with developmental temperature, where higher temperatures during development led to smaller individuals (Spearman’s *ρ* = –0.566, p < 0.001). These effects were present in both male and female animals, however, females had significantly higher thermal tolerance and larger body size compared to males (CT*_max_* females: 28.4 °C *±*1.21 °C, males: 27.7 °C *±*1.21 °C, length females: 744 µm *±*69 µm, males: 694 µm *±*44 µm, p *<* 0.001 for both).

In addition to the correlation of developmental temperature, there was a negative correlation between prosome length and CT*_max_* in wild animals where smaller individuals had significantly higher thermal tolerance (p *<* 0.001, Pearson’ r = –0.229, Fig. 4A). While this result was likely driven by the aforementioned relation of developmental temperature and length, the negative trend was also present, though weaker, when looking only at the warm treatment common gardened animals (p = 0.038, Fig. 4B); no effect was observed within cold common gardened treatments (p = 0.108, Fig. 4B).

**Figure 4:**
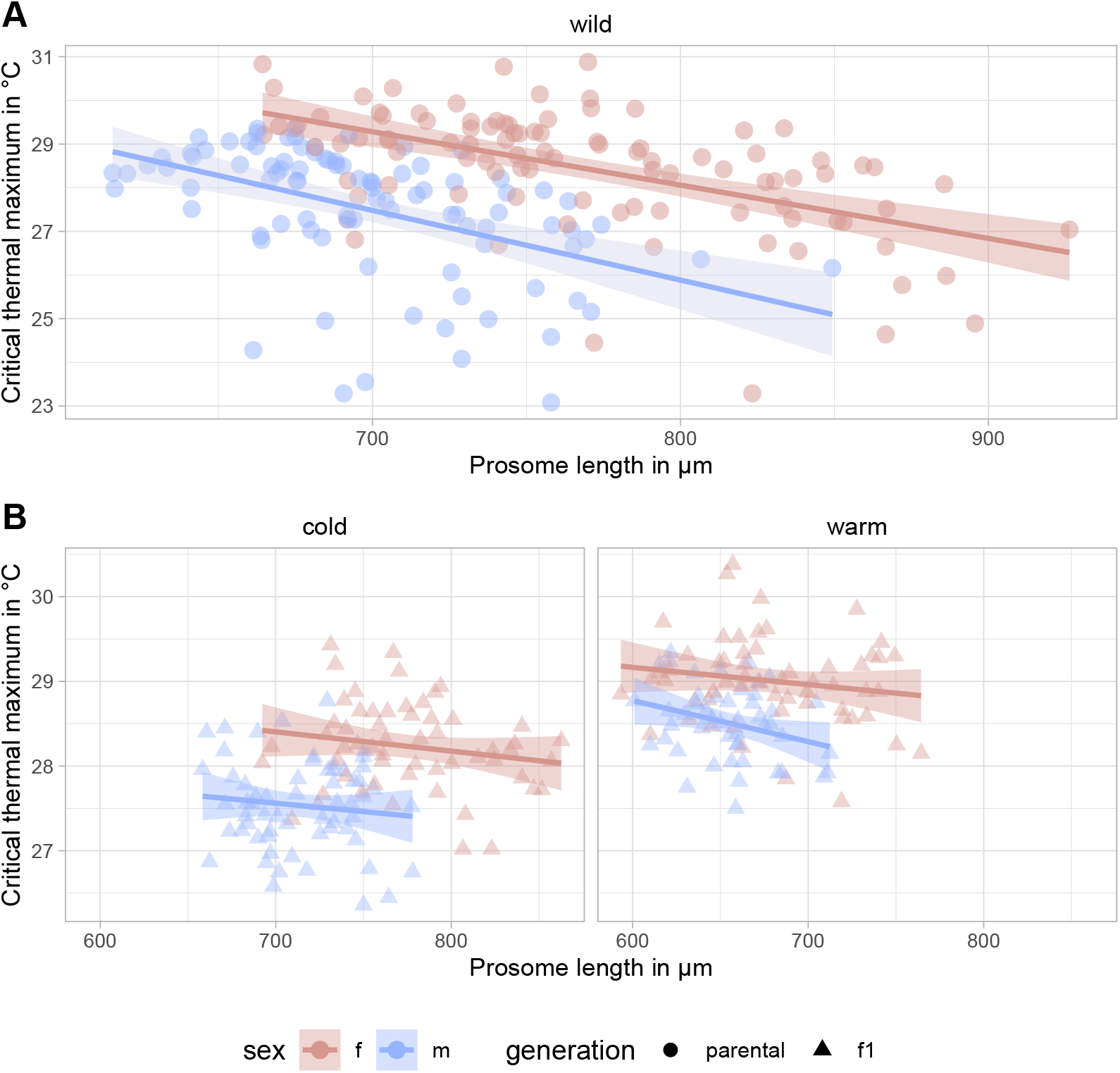
Correlation between prosome length and CT*_max_* in *A. hudsonica*, (A) Parental generation, length has a significant effect on CT*_max_* (p < 0.001, R^2^ = 0.35) (B) F1 generation; cold treatment: no significant effect of length on CT*_max_* (p = 0.108, R^2^ = 0.323); warm treatment: a significant effect of length on CT*_max_* (p = 0.038, R^2^ = 0.242). Linear regression per sex with 95% confidence interval.

## 4. Discussion

Seasonal fluctuations have strong physiological effects on organisms occupying these variable conditions. We hypothesized that the thermal tolerance of copepods would closely mirror their developmental temperature and that both plastic and genetic mechanisms would contribute to the physiological change observed. As predicted, the CT*_max_* of *A. hudsonica* increased in parallel with environmental temperature. However, under common garden conditions collections showed similar levels of plasticity and converged on common thermal tolerances and body sizes, indicating that phenotypic differences between collection times were driven by plasticity with no evidence for rapid adaptation. Together, these results indicate that *A. hudsonica* has substantial phenotypic plasticity to rapidly acclimate to large changes in external temperature.

### 4.1. Seasonal variation in thermal tolerance

For both marine and terrestrial ectotherms, the ability to rapidly acclimate to changes in environmental temperature is common and adaptive (Gunderson and Stillman, 2015). In coastal marine organisms, particularly those from temperate environments, the presence of thermal plasticity is essential as shallow waters tend have high variance in their temperatures, requiring rapid phenotypic responses (Reusch, 2014). This is particularly true for intertidal copepods which can experience daily temperature changes of nearly 10°C (Leong et al., 2017) and therefore have high thermal tolerance and plasticity (Healy et al., 2019). Further, across copepods there is near universal presence of thermal plasticity that is dependent on environmental temperature (Sasaki and Dam, 2021). For *Acartia* copepods specifically, *A. tonsa* and *A. hudsonica* from the east coast of the United States are typically plastic in their thermal tolerance (Sasaki and Dam, 2020) and laboratory studies on *A. tonsa* show large effects of acclimation on thermal tolerance (Sunar and Kir, 2021). Thus, plasticity plays an important role in enabling most marine copepods to tolerate environmental temperature fluctuations.

While the Baltic Sea experiences only small wind-driven and irregular tides, the variation in water temperature at our collection site is nevertheless high, ranging from 5.65 °C to 20.70 °C during the study period. This high and relatively predictable variation likely favors the evolution of plasticity observed in the population (Bitter et al., 2021). Indeed, previous work has shown that *Acartia tonsa* from less variable low latitude thermal environments harbor lower levels of phenotypic plasticity than those from more variable high latitude sites (Sasaki and Dam, 2020), consistent with the latitudinal hypothesis of plasticity (Janzen, 1967; Ghalambor et al., 2006). Similarly, Sasaki and Dam (2020) found that seasonal variation in thermal LD50 of *A. hudsonica* from the east coast of North America was driven by plasticity rather than adaptation. Therefore, the plasticity that *A. hudsonica* harbors to respond to changing temperature across the season is likely adaptive in this environment and is present across multiple populations.

The lack of seasonal adaptation is in contrast to the evidence of this phenomenon in other systems including both terrestrial *D. melanogaster* (Johnson et al., 2023) and the sister species to *A. hudsonica*, *A. tonsa* (Sasaki and Dam, 2020). For *A. tonsa*, Sasaki and Dam (2020) found that collections from different time points differed in their plasticity. It is unclear exactly what drove the seasonal patterns in *A. tonsa*, but it may be due to the presence of cryptic lineages emerging or surviving at different temperatures. The comparison to *D. melanogaster* is also interesting as terrestrial organisms typically have a better ability than marine organisms to buffer their thermal environment via behavioral mechanisms, known as the Bogert effect (Bogert, 1949). *Acartia* copepods have enormous populations sizes that are regularly in the hundreds of individuals per cubic meter of water (Möllmann, 2002). Therefore, the efficiency of selection may be similar between *Acartia* and *Drosophila*. Given this, one might expect similar signals of seasonal adaptation in our system relative to *Drosophila*; we observed no evidence to support this expectation.

There are a number of possible explanations for the lack of seasonal adaptation in our study. First, *A. hudsonica* may have sufficient plasticity to respond to the seasonal thermal environment. Given the Bogert effect, environmental temperature is directly experienced to a greater degree in marine systems and selection for plasticity may be strong and result in a highly flexible phenotype. Alternatively, there may be no heritable genetic variation for thermal tolerance in this population. However, populations of *A. hudsonica* from North America can rapidly evolve to elevated temperatures under laboratory conditions (deMayo et al., in press) and there is evidence for local adaptation to temperature along latitudinal gradients in the sister species, *Acartia tonsa* (Sasaki and Dam, 2019). Therefore, it is likely that heritable variation in thermal tolerance is also present in this species and population. An alternative explanation is that the phenotypes of focus were not under selection or sensitive enough to capture any adaptive responses.

The temperatures at the collection site did not approach the critical thermal limit measured in the lab (maximum SST during sampling: 20.70 °C). Therefore, it is unlikely that CT*_max_* was directly under selection. It is possible that an alternative phenotype would show a seasonal adaptive response as has been observed in other systems (Hairston and Dillon, 1990; de Villemereuil et al., 2020). Finally, much of the evidence for seasonal adaptation in *D. melanogaster* has been found at the genomic level (Johnson et al., 2023). Given this, our populations may similarly be experiencing fluctuating selection that would be detectable using genomic approaches.

While we observed high levels of plasticity for CT*_max_*, under future temperature conditions it is unlikely that most ectotherms have sufficient plasticity to respond to temperature changes without adaptive responses (Gunderson and Stillman, 2015). Indeed, DeMayo et al showed that under warming conditions, plasticity alone is insufficient to maintain high population fitness in *A. hudsonica* (deMayo et al., in press). However, the species can rapidly adapt after just four generations to recover fitness levels. Therefore, it is likely that both plasticity and adaptation will be required to tolerate future environmental conditions and more work is needed to understand the relative contribution of each to overall resilience.

### 4.2. Plasticity in body size and potential potential impacts of warming

As temperatures increased, body size decreased in *A. hudsonica* (Fig. 1), a common phenomenon across ectotherms known as the temperature-size rule (Angilletta and Dunham, 2003; Rubalcaba and Olalla-Tárraga, 2020). This concept applies to copepods (Escribano and McLaren, 1992; Viitasalo et al., 1995), including those from tropical (Ortega-Mayagoitia et al., 2018) and temperate environments (Riccardi and Mariotto, 2000). However, other factors such as phytoplankton density may affect the body size of individuals in the wild (Deevey, 1964, 1966), though this would not have affected the common garden animals in our study. The reduction in body size in response to increasing temperature may be driven by the disproportionate increase in respiration and metabolism relative to ingestion and assimilation of nutrients (Lehman, 1988), leading to lower overall energy available for growth and therefore a smaller body size. Further, there may be a trade-off that favors smaller individuals at high temperatures; reproductive efficiency, the ratio of egg production and respiration, is maximized at smaller body sizes and therefore may be adaptive in warmer temperatures. Similarly, at higher temperatures oxygen availability (aerobic scope) may be decreased in larger individuals relative to smaller individuals, favoring smaller body sizes (Rubalcaba et al., 2020). Alternatively, in cold temperatures growth periods may be prolonged while the growth rate remains relatively stable, leading to larger individuals under cold conditions (Vidal, 1980).

Regardless of the mechanism, as temperatures warm due to anthropogenic causes, decreases in body size may affect ecosystem interactions. This is particularly true in the Baltic Sea where the heating rate is around three times higher than the ocean average due to its unique topography (Reusch et al., 2018; Szymczycha et al., 2019; Dutheil et al., 2022). A size reduction in *A. hudsonica*, or other prey organisms, might impact higher trophic level predators, for example by requiring the consumption of more individuals to maintain the same amount of nutrient intake (Garzke et al., 2015). If abundance does not increase with decreasing body size, nutrient availability may be reduced for consumers who will also require increased energy needs under higher temperature (Brown et al., 2004). Further, Garzke et al. (2015) observed that large copepods from colder temperatures clear algae biomass more efficiently than smaller individuals, exerting top-down control on phytoplankton. This in turn suggests that smaller copepods are less efficient grazers, with less control over the planktonic community. As copepods are important grazers on large phytoplankton and microzooplankton (Sommer et al., 2003; Armengol et al., 2017), less efficient grazing or a shift to different prey size classes might have unforeseen cascading effects across ecosystems. Finally, the correlation between body size and CT*_max_* (Fig. 4) suggests that body size itself may influence the thermal tolerance of an individual with small body size being of advantage in warm environments. While the effect was weak in comparison with the effects of developmental temperature and the mechanistic link between body size and thermal tolerance remains unknown, this relationship could potentially be used to predict an individual’s thermal tolerance. This may be of interest when analyzing historical samples where body size measurements from the same location across years could enable predictions about past thermal tolerance and environmental temperatures. More work would be needed to develop these predictions.

### 4.3. Outliers and mixed species

The outliers in collection 2 under cold conditions, characterized by unexpectedly low thermal tolerance and small body sizes, were most likely the result of an impending culture collapse. This was potentially caused by poor food quality during that period due to ciliate and bacteria growing in the algae cultures used for feeding. The negative effects of ciliates on copepod fitness are well described with effects ranging from decreased egg production (Burris and Dam, 2014) to increased adult mortality (Visse, 2007). The species identity of the ciliates in this study could not be determined. However, ciliate peak abundances correlated with culture collapse, and after the establishment of more frequent water changes, the cultures improved. Since the survival and fitness of the animals was clearly affected by external factors unrelated to the experiment, length and thermal tolerance measurements for the cold treatment in collection 2 should be treated with caution and were therefore excluded from parts of the analysis.

In the wild collection 5, five individuals showed a CT*_max_* above 33 °C, which was well beyond the distribution of values for any other collection (Fig. S.5). In the F1 generation, the warm treatment of collection 4 and 5 and the cold treatment of collection 5 also had similar high performing individuals (11, 40, and 2 individuals, respectively) (Fig. S.5). These extreme outliers suggested a mixed species composition, which was confirmed by genotyping individuals of each collection as well as high thermal outliers (see Supplement *Mixed species*). High thermal outlier individuals were *A. tonsa* without exception (Table S.1, Fig. S.6). The increased thermal tolerance for *A. tonsa* relative to *A. hudsonica* is consistent with other studies. On the east coast of North America, *A. hudsonica* dominates plankton communities early in the year when water temperatures are low and is replaced by *A. tonsa* as temperatures increase (Borkman et al., 2018; Sullivan and McManus, 1986). This pattern appears to be similar for copepod communities in Kiel Bight as well. Since *A. tonsa* was present only in later collections and just as individual outliers, this paper could not determine how thermal tolerance of *A. tonsa* changes within a season. However, we would hypothesize that *A. tonsa* follows an overall similar trend than shown for *A. hudsonica* with thermal tolerances shifted towards warmer temperatures. Additional experiments would be required to test this hypothesis.

### 4.4. First record of A. hudsonica in the Baltic Sea

From our literature and database review, the molecular barcoding in this study is the first record of *A. hudsonica* in the Baltic Sea. There are two possible explanations for this novel species presence. First, *A. hudsonica* may have recently invaded the Baltic Sea. In the North Sea, *A. omori* – a species that co-occurs with *A. hudsonica* off the Japanese coast (Ueda, 1987) – was first described by Seuront (2005) in the early 2000. The successful invasion of *A. omori* suggests that *A. hudsonica* could similarly have been introduced to the North Sea. As the North Sea and Baltic Sea are connected, it is possible that *A. hudsonica* then moved to the Baltic Sea. Alternatively, there is ample shipping in the region and an independent local introduction could have occurred.

Secondly, it cannot be ruled out that *A. hudsonica* has historically been misidentified in the Baltic Sea. Copepods from the genus *Acartia* have a record of mis-identifications in databases and are difficult to distinguish morphologically (Figueroa et al., 2020). Instead, phylogenetic methods are required for accurate species identification. Before the 1970’s, *A. hudsonica* was a subspecies of *A. clausi*, but is now considered its own species (Bradford, 1976; Ueda, 1986). Given this, it is possible that *A. hudsonica* is native to the Baltic Sea but has been, and still is, commonly identified as *A. clausi*.

## 5. Conclusions

Here we showed that *Acartia hudsonica* has high phenotypic plasticity in response to changing temperature within a single year. We found that thermal tolerance closely tracks environmental temperature, indicating that *A. hudsonica* has capacity to tolerate increasing temperatures that fall within the current range experienced in nature. However, the observed decrease in body size suggests that nutrient availability and ecosystem functioning could be impacted if temperatures consistently increase with no change in copepod abundance. By focusing on the relative impacts of plasticity and adaptation to population responses to temperature change we can begin to understand the resilience populations and ecosystems to ongoing global change.

## Acknowledgement

We thank Catriona Clemmesen-Bockelmann for her support during the project and Tobias Strickmann for collecting the samples. We thank Fabian Wendt and Diana Gill for their guidance in animal cultivation and lab work. Further, we thank Gianina Consing, Sheena Chung, Myria Schröder and Friederike Gronwald for assisting with experimental work and lab work. This project was partially funded by the German Research Foundation BR6839/1-1 (to R.B).

## Data accessibility

The data set and scripts can be found on https://github.com/HahnAlexandra/ Plasticity_Acartia_hudsonica. The individual sequences are uploaded on NCBI GenBank https://www.ncbi.nlm.nih.gov/genbank/, find individual accession numbers in Table S.1. SST data from Kiel Fjord will be made available on https://www.pangaea.de by our collaborator Claas Hiebenthal.

