## Supplemental Information for "Phenotypic Plasticity Drives Seasonal Thermal Tolerance in a Baltic Copepod"

### Supplements

#### *Mixed species*

For each collection, random individuals were sequenced to verify species identity (see Table S.1). For the first two time points, phylogenetic mapping of mtCOI sequences indicated the uniform presence of *A. hudsonica*. In later collections, *A. hudsonica* co-occurred with *A. tonsa*, requiring further sequencing efforts. Collections 4 and 5 included individuals with $CT_{max}$ measurements up to 5 °C higher than the highest non-outlier measurements. To confirm outlier identities in collections 4 and 5, 38 additional samples, including outliers as well as randomly sampled non-outlier controls, were genotyped. Three samples failed to sequence and were dropped from the analysis.

Phylogenetic analysis showed that high thermal outliers were *A. tonsa* clade X, while lower reference samples matched to *A. hudsonica* (Fig. S.6). The experimental animals formed two clearly divided clusters along a thermal performance gradient (Fig. S.7). Since the majority of experimental animals between different collections belonged to one species, only *A. hudsonica* was considered for the main analysis.

### *Supplementary Tables*

Table S.1: Overview of individuals sequenced including NCBI genbank accessions, Random individuals were taken from the original cultures, outliers were high-performing experimental individuals and non high-performing controls.

| Origin | ID | Species | NCBI | Origin | ID | Species | NCBI |
|---|---|---|---|---|---|---|---|
| **Random individuals** | | | | **Outliers** | | | |
| Collection 1 wild | Col1_1 | A. hudsonica | OR345065 | Collection 4 wild | 220630L10 | failed | - |
| Collection 1 wild | Col1_2 | A. hudsonica | OR345066 | Collection 4 wild | 220629L4 | failed | - |
| Collection 1 wild | Col1_3 | A. hudsonica | OR345067 | Collection 4 wild | 220630R6 | A. hudsonica | OR345092 |
| Collection 1 wild | Col1_4 | A. hudsonica | OR345068 | Collection 4 wild | 220629R9 | A. hudsonica | OR345091 |
| Collection 1 wild | Col1_5 | A. hudsonica | OR345069 | Collection 4 warm | 220714R6 | A. tonsa | OR345098 |
| Collection 1 wild | Col1_6 | A. hudsonica | OR345070 | Collection 4 warm | 220714R3 | A. hudsonica | OR345096 |
| Collection 1 wild | Col1_7 | A. hudsonica | OR345071 | Collection 4 warm | 220714R5 | A. hudsonica | OR345097 |
| Collection 1 wild | Col1_8 | A. hudsonica | OR345072 | Collection 4 warm | 220715L9 | A. hudsonica | OR345103 |
| Collection 1 wild | Col1_9 | A. hudsonica | OR345073 | Collection 4 warm | 220714L1 | A. tonsa | OR345093 |
| Collection 1 wild | Col1_10 | A. hudsonica | OR345074 | Collection 4 warm | 220714L6 | A. tonsa | OR345094 |
| Collection 1 wild | Col1_11 | failed | - | Collection 4 warm | 220714L9 | A. tonsa | OR345095 |
| Collection 1 wild | Col1_12 | A. hudsonica | OR345075 | Collection 4 warm | 220714R8 | A. tonsa | OR345099 |
| Collection 1 wild | Col1_13 | A. hudsonica | OR345076 | Collection 4 warm | 220714R9 | A. tonsa | OR345100 |
| Collection 1 wild | Col1_14 | failed | - | Collection 4 warm | 220715R6 | A. tonsa | OR345105 |
| Collection 1 wild | Col1_15 | failed | - | Collection 4 warm | 220715R7 | A. tonsa | OR345106 |
| Collection 2 wild | Col2_1 | failed | - | Collection 4 warm | 220715R10 | A. tonsa | OR345104 |
| Collection 2 wild | Col2_4 | failed | - | Collection 4 warm | 220715L4 | A. tonsa | OR345101 |
| Collection 2 wild | Col2_6 | failed | - | Collection 4 warm | 220715L5 | A. tonsa | OR345102 |
| Collection 2 wild | Col2_7 | failed | - | Collection 5 wild | 220724L6 | A. hudsonica | OR345116 |
| Collection 2 wild | Col2_8 | A. hudsonica | OR345077 | Collection 5 wild | 220721L9 | failed | - |
| Collection 2 wild | Col2_13 | failed | - | Collection 5 wild | 220721L4 | A. hudsonica | OR345108 |
| Collection 2 wild | Col2_14 | A. hudsonica | OR345078 | Collection 5 wild | 220722L3 | A. hudsonica | OR345111 |
| Collection 2 wild | Col2_15 | A. hudsonica | OR345079 | Collection 5 wild | 220721R5 | A. hudsonica | OR345110 |
| Collection 2 wild | Col2_16 | failed | - | Collection 5 wild | 220721L8 | A. hudsonica | OR345109 |
| Collection 3 wild | Col3_3 | failed | - | Collection 5 wild | 220721L1 | A. tonsa | OR345107 |
| Collection 3 wild | Col3_4 | A. hudsonica | OR345080 | Collection 5 wild | 220722L8 | A. tonsa | OR345112 |
| Collection 3 wild | Col3_7 | A. hudsonica | OR345081 | Collection 5 wild | 220722R2 | A. tonsa | OR345113 |
| Collection 3 wild | Col3_9 | A. hudsonica | OR345082 | Collection 5 wild | 220722R5 | A. tonsa | OR345114 |
| Collection 4 wild | Col4_1 | failed | | Collection 5 wild | 220722R8 | A. tonsa | OR345115 |
| Collection 4 wild | Col4_2 | A. hudsonica | OR345083 | Collection 5 cold | 220823L4 | A. hudsonica | OR345122 |
| Collection 4 wild | Col4_3 | A. hudsonica | OR345084 | Collection 5 cold | 220823R5 | A. hudsonica | OR345124 |
| Collection 4 wild | Col4_5 | A. hudsonica | OR345085 | Collection 5 cold | 220823R10 | A. tonsa | OR345123 |
| Collection 4 wild | Col4_6 | A. hudsonica | OR345086 | Collection 5 cold | 220824L8 | A. tonsa | OR345125 |
| Collection 4 wild | Col4_7 | A. hudsonica | OR345087 | Collection 5 warm | 220804L2 | A. tonsa | OR345117 |
| Collection 4 wild | Col4_9 | failed | - | Collection 5 warm | 220804L6 | A. tonsa | OR345118 |
| Collection 5 wild | Col5_1 | failed | - | Collection 5 warm | 220804R7 | A. tonsa | OR345119 |
| Collection 5 wild | Col5_2 | failed | - | Collection 5 warm | 220805L8 | A. tonsa | OR345120 |
| Collection 5 wild | Col5_3 | failed | - | Collection 5 warm | 220805L9 | A. tonsa | OR345121 |
| Collection 5 wild | Col5_4 | A. hudsonica | OR345088 | | | | |
| Collection 5 wild | Col5_5 | A. hudsonica | OR345089 | | | | |
| Collection 5 wild | Col5_6 | failed | - | | | | |
| Collection 5 wild | Col5_8 | failed | - | | | | |
| Collection 5 wild | Col5_9 | A. hudsonica | OR345090 | | | | |

Table S.2: Overview over sampling and assay dates.

| **Collection name** | **Collection date** | **Wild** | **F1 warm** | **F1 cold** |
|---|---|---|---|---|
| Collection 1 | 06 Apr 2022 | 07/08 Apr | 26/27 Apr | 10/11 May |
| Collection 2 | 16 May 2022 | 17/18 May | 08/10 May | 23/28 Jun |
| Collection 3 | 13 Jun 2022 | 15/16 Jun | - | - |
| Collection 4 | 27 Jun 2022 | 29/30 Jun | 14/15 Jul | 08/09 Aug |
| Collection 5 | 19 Jul 2022 | 21/22 Jul | 04/05 Aug | 23/24 Aug |

Table S.3: ANOVA type III results for the regression relating $CT_{max}$ to collection, treatment, sex, and length, model for *A. hudsonica.*

| | **d.f.** | **Sum. Sq.** | **F-value** | **Pr (>F)** |
|---|---|---|---|---|
| **$CT_{max}$** | | | | |
| collection | 3 | 194.659 | 141.102 | $<2e^{-16}$ |
| treatment | 2 | 121.735 | 132.363 | $<2e^{-16}$ |
| collection x treatment | 5 | 178.759 | 77.747 | $<2e^{-16}$ |
| sex | 1 | 36.503 | 79.380 | $<2e^{-16}$ |
| length | 1 | 0.578 | 1.256 | 0.2631 |
| **Length** | | | | |
| collection | 3 | 273014 | 67.019 | $<2e^{-16}$ |
| treatment | 2 | 197310 | 72.654 | $<2e^{-16}$ |
| collection x treatment | 5 | 195872 | 28.850 | $<2e^{-16}$ |
| sex | 1 | 305356 | 224.876 | $<2e^{-16}$ |

Table S.4: ANOVA type III results for the effects of developmental temperature on length and thermal tolerance.

| | **d.f.** | **Sum. Sq.** | **F-value** | **Pr (>F)** |
|---|---|---|---|---|
| **$CT_{max}$** | | | | |
| developmental temp | 1 | 306.7 | 442.197 | $<2e^{-16}$ |
| sex | 1 | 35.5 | 51.141 | $4.227e^{-12}$ |
| ***Length*** | | | | |
| developmental temp | 1 | 570342 | 289.09 | $<2e^{-16}$ |
| sex | 1 | 252079 | 127.77 | $<2e^{-16}$ |

*Supplementary Figures*

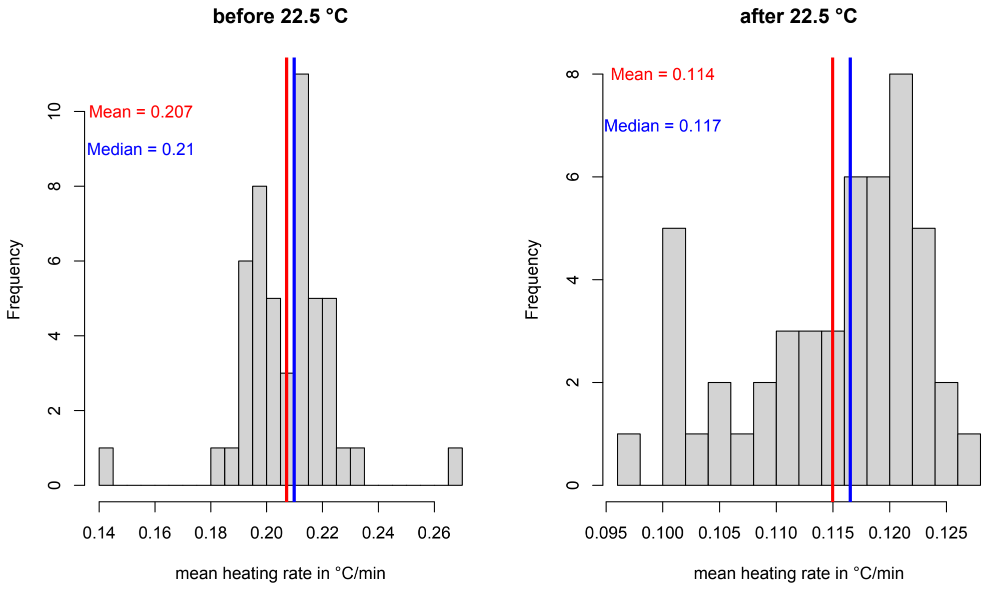


Figure S.1: Mean ramping rates during the experiment before and after unplugging the second heater, vertical lines indicate mean and median of all trials.

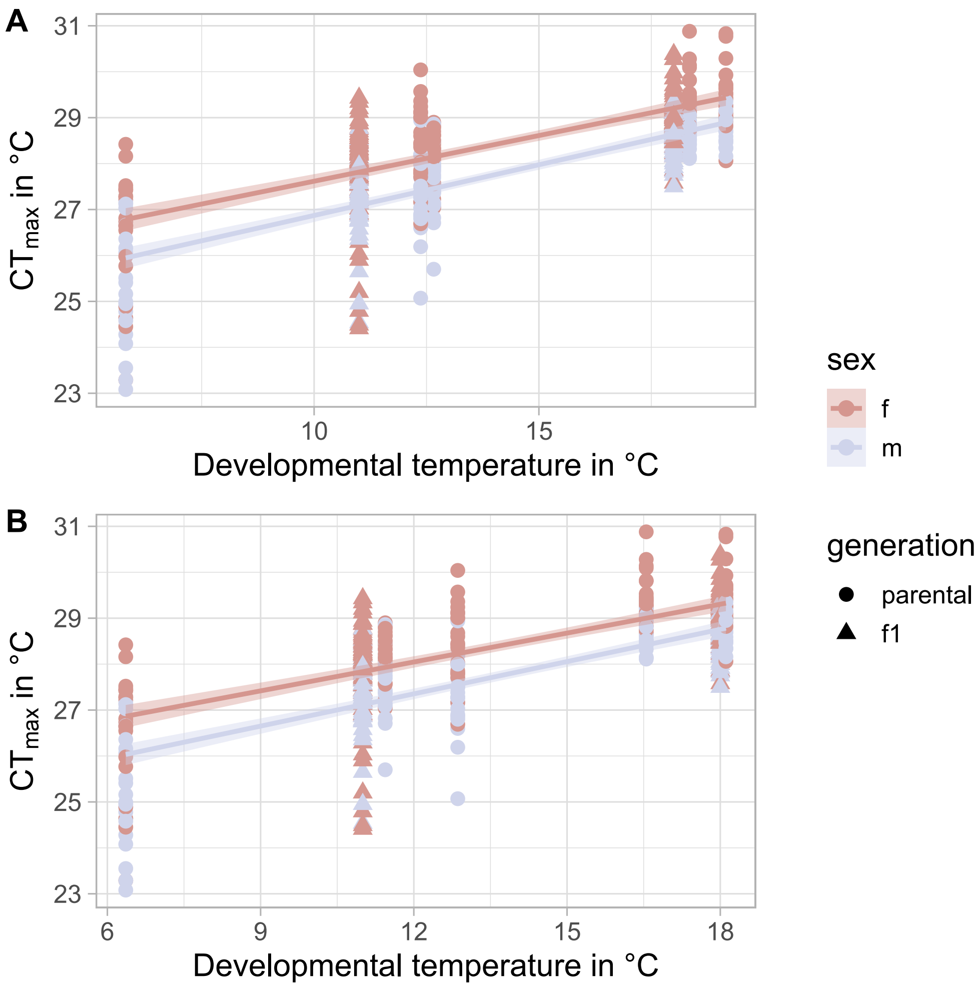


Figure S.2: Correlation between $CT_{max}$ and developmental temperature for all experimental individuals, A: mean SST on sampling day, Spearman's $\rho = 0.711$, p $< 0.001$; B: mean SST for the 2-week period prior to sampling, Spearman's $\rho = 0.689$, p $< 0.001$.

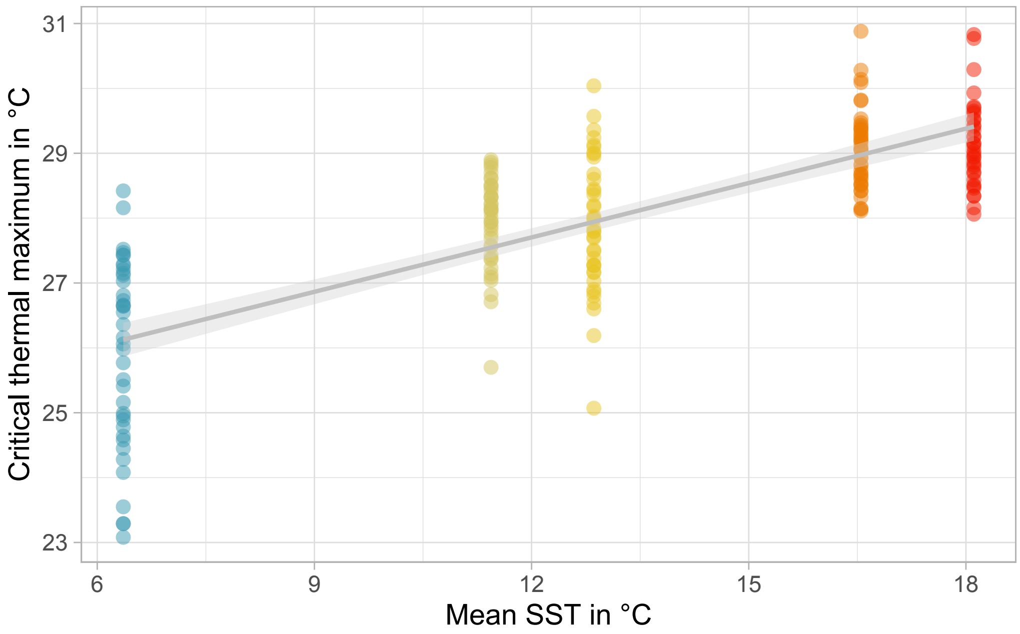


Figure S.3: Positive correlation between mean SST (averaged over the two weeks prior to sampling) and wild $CT_{max}$, Spearman's $\rho = 0.758$, $p < 0.001$, $R^2 = 0.585$, linear regression with 95 % confidence interval.

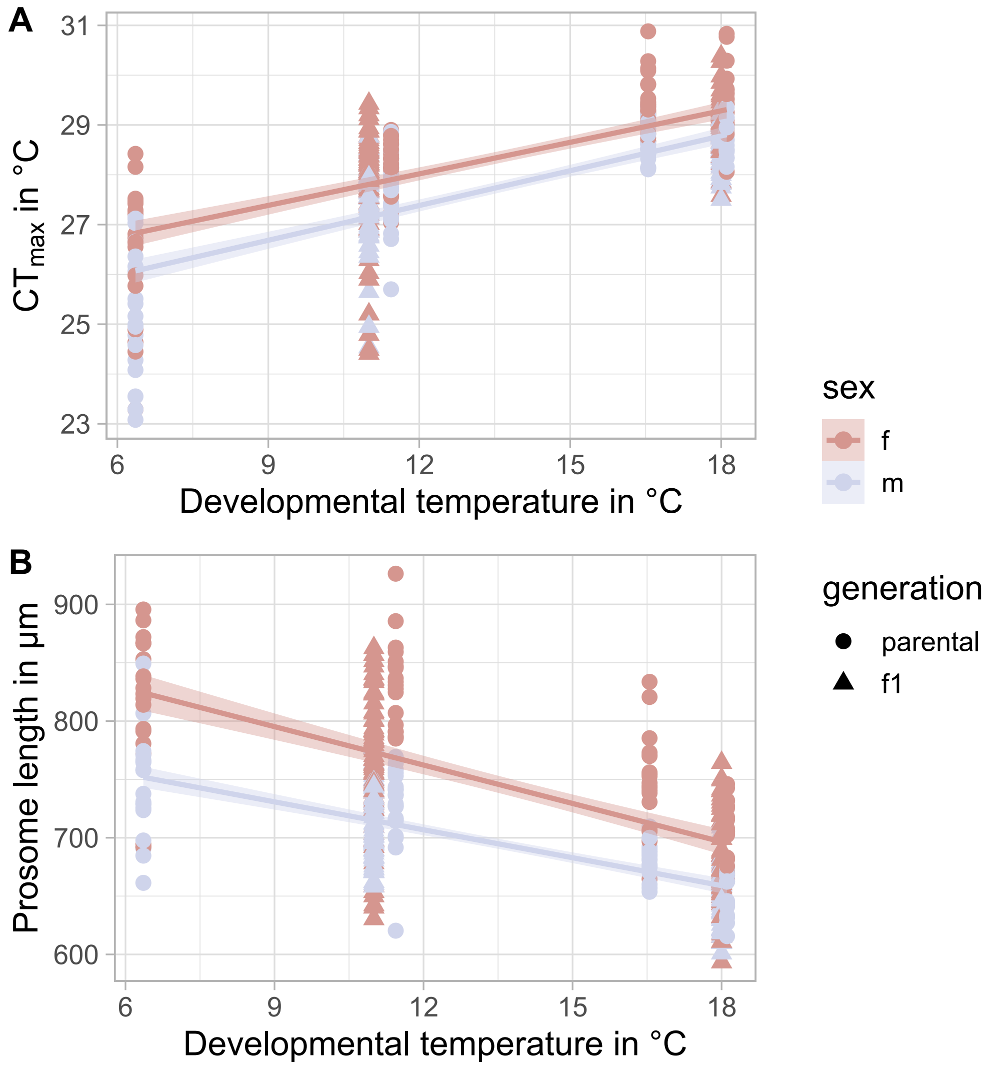


Figure S.4: Effects of developmental temperature on A: $CT_{max}$ (developmental temperature: $p < 0.001$, sex: $p < 0.001$), $R^2 = 0.557$, and B: prosome length (developmental temperature: $p < 0.001$, sex: $p < 0.001$), $R^2 = 0.510$; Linear regression per sex with 95% confidence interval for all treatments in collections 1,2,4 and 5.

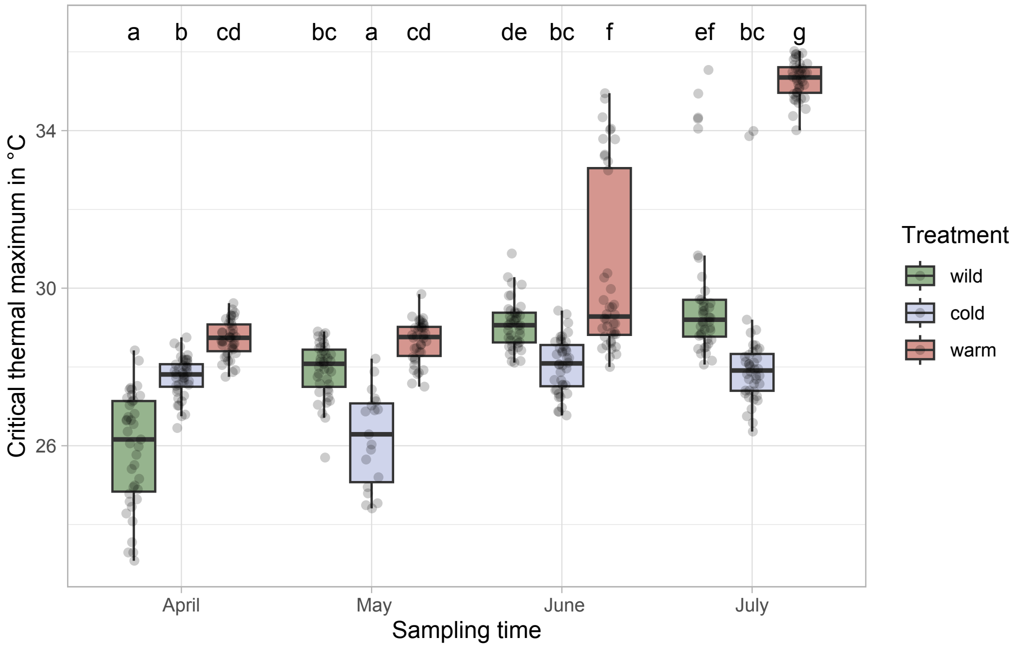


Figure S.5: Critical thermal maxima for all experimental individuals, *A. hudsonica* and *A. tonsa*. Colours of boxes correspond to treatment. Compact letters based on weighted means per group, same letters indicate no significant difference between measurements.

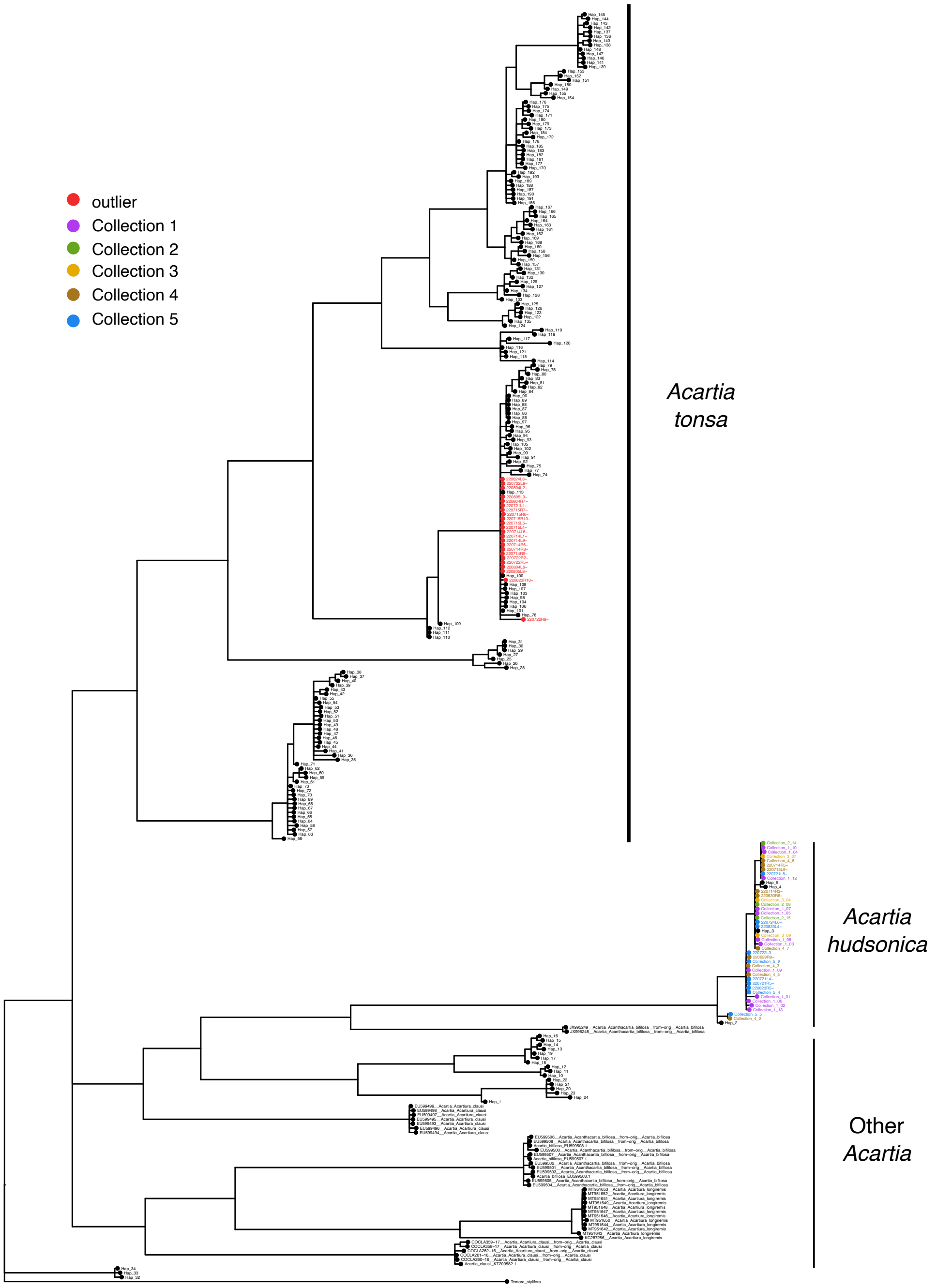


Figure S.6: Phylogenetic tree of all genotyped individuals mapped to sequences of known species identity. Based on mtCOI and inferred by Bayesian methods.

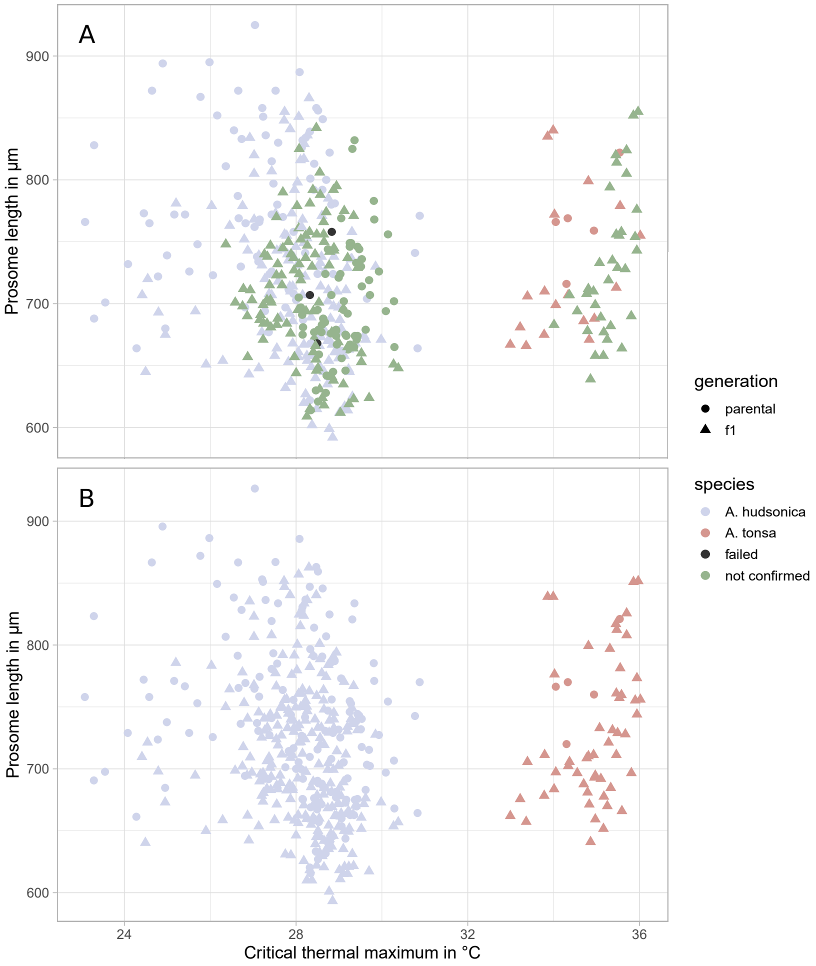


Figure S.7: Species identity shown by length and thermal tolerance; A: confirmed species identity through genotyping, B: assumed species identity based on proximity to confirmed individuals, all individuals in the low thermal tolerance cluster are assumed to be *A. hudsonica*, all individuals in the high thermal tolerance cluster are assumed to be *A. tonsa*.

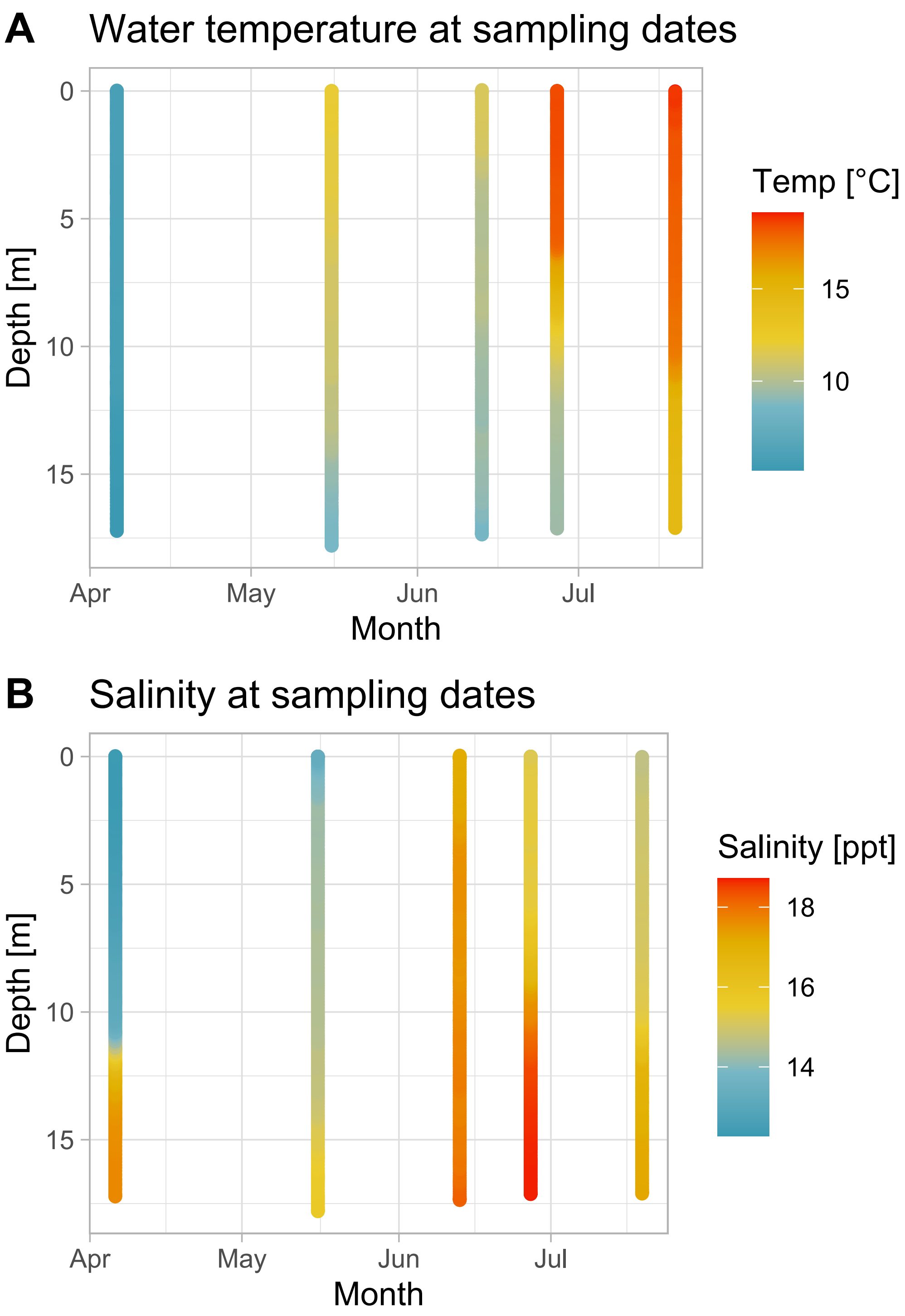


Figure S.8: Temperature and salinity data recorded by *RV Polarfuchs* on sampling dates; A: temperature data, B: salinity data.

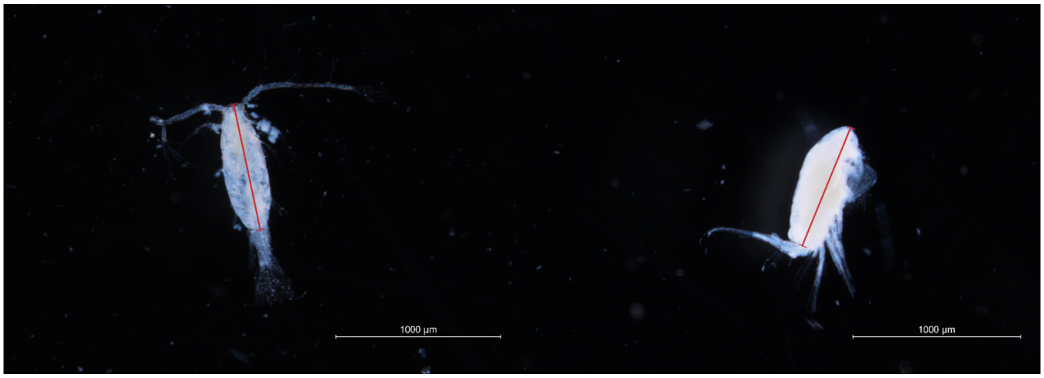


Figure S.9: Images of experimental copepods, red bars indicate the measured prosome length, left: male in top view, right: female in side view.
